# The *C. elegans* SET-2 histone methyltransferase maintains germline fate by preventing progressive transcriptomic deregulation across generations

**DOI:** 10.1101/583799

**Authors:** Valérie J. Robert, Andrew K. Knutson, Andreas Rechtsteiner, Gaël Yvert, Susan Strome, Francesca Palladino

## Abstract

Chromatin factors contribute to germline maintenance by preserving a germline-appropriate transcriptional program. In the absence of the conserved histone H3 Lys4 (H3K4) methyltransferase SET-2, *C. elegans* germ cells progressively lose their identity over generations, leading to sterility. How this transgenerational loss of fertility results from the absence of SET-2 is unknown. Here we performed expression profiling across generations on germlines from mutant animals lacking SET-2 activity. We found that gene deregulation occurred in 2 steps: a priming step in early generations progressing to loss of fertility in later generations. By performing Within-Class Analysis (WCA), a derivative of Principal Component Analysis, we identified transcriptional signatures associated with SET-2 inactivation, both at the priming step and later on during loss of fertility. Further analysis showed that repression of germline genes, derepression of somatic programs, and X-chromosome desilencing through interference with PRC2-dependent repression, are priming events driving loss of germline identity in the absence of SET-2. Decreasing expression of identified priming genes, including the C/EBP homologue *cebp-1* and TGF-β pathway components, was sufficient to delay the onset of sterility, suggesting that they individually contribute to the loss of germ cell fate. Altogether, our findings illustrate how the loss of a chromatin regulator at one generation can progressively deregulate multiple transcriptional and signaling programs, ultimately leading to loss of appropriate cell fate.

## Introduction

Maintenance of cell fate requires activating lineage-appropriate genes and repressing lineage-inappropriate genes. This process has been extensively studied *in vitro* in cultured pluripotent cells (iPSCs) and shown to rely on appropriate signaling, transcription factor binding, and chromatin states (Lessard and Crabtree 2010; Orkin and Hochedlinger 2011). Much less is known about how cell identity is maintained in an intact tissue. The preservation of germ-cell identity is essential for germline survival and the passage of genetic information from generation to generation. In *C. elegans*, germ-cell identity is maintained by translational regulators, germ granules (called P granules in *C. elegans*), and chromatin factors including Polycomb Repressive Complex 2 (PRC2), composed of MES-2, MES-3, and MES-6 (Ciosk, DePalma, and Priess 2006; Updike et al. 2014; Patel et al. 2012; Tursun et al. 2011; Robert et al. 2014), and the H3K36 methyltransferase MES-4, a homolog of the vertebrate NSD proteins (Bender et al. 2006; Fong et al. 2002; Bender et al. 2004). PRC2 and MES-4 cooperate to produce a functional germline through repression of the X chromosome and establishment of gene expression programs appropriate for germ cells (Bender et al. 2004; Xu, Fong, and Strome 2001; Gaydos et al. 2012). In the adult germline, PRC2 buffers germ cells against reprogramming: forced expression of a master regulatory protein in the absence of PRC2 components leads to conversion of germ cells into somatic cell types (Patel et al., 2012). Increased GLP-1/Notch signaling in the germline antagonizes PRC2-mediated silencing of somatic genes to promote reprogramming (Seelk et al. 2016).

Compromising germline H3K4 methylation levels through inactivation of the conserved SET-2/SET1 histone methyltransferase or the H3K4 demethylase SPR-5/LSD1 also results in loss of germline identity (Robert et al., 2014; Xiao et al., 2011; Katz et al., 2009). Interestingly, while SET-2 inactivation results in an immediate decrease in H3K4 methylation in the germline, loss of germline identity is only observed under stress conditions and takes place over multiple generations, culminating in sterility and the conversion of germ cells into somatic cell types (Xiao et al. 2011; Robert et al. 2014).

Expression profiling of germlines lacking P-granule components, PRC2 or MES-4 (Gaydos et al. 2012; Knutson et al. 2017; Campbell and Updike 2015), or undergoing forced reprogramming in the context of increased GLP-1/Notch signaling (Seelk et al. 2016), has shown that loss of germ-cell identity is associated with expression of somatic genes. However, these approaches could not distinguish between genes whose misexpression causes versus is a consequence of a perturbed germline transcriptional program and sterility. The progressive loss of germline identity in germlines lacking SET-2 activity provides the opportunity to compare transcriptional profiles from early-generation mutant germlines in which fertility is not compromised and late-generation fertile and sterile animals. In this study, transgenerational transcriptomic data was analyzed using Within-Class Analysis (WCA), a multivariable analysis strategy, on paired data sets. This approach, which removes variability between lineages, allowed the identification of a subset of genes consistently misregulated between individual lineages, both in early-generation fertile animals and in later-generation sterile animals. Our approach emphasizes that downregulation of germline gene expression, upregulation of somatic gene expression and X-chromosome desilencing are priming events of loss of germline identity in the absence of SET-2. Interestingly, our approach also revealed that components of the conserved TGF-beta signaling pathway, including receptors and downstream effectors, are consistently upregulated during loss of germline identity, suggesting a novel function for TGF– β signaling in maintaining germline identity in *C. elegans*.

## Results

### Loss of germline identity in *set-2* mutants correlates with progressive and widespread transcriptional deregulation

We determined transcriptome changes that accompany loss of germline identity and may underlie the sterility of late-generation *set-2* mutant animals. From *set-2/+* heterozygous animals grown at 25°C (P0 generation), we derived *set-2/set-2* homozygous animals that we grew for 1 additional generation (F2 - fertile) and for a total of 4 generations (F4 - both fertile and sterile) (Figure 1A). Germline dissections and RNA sequencing were carried out on 3 separate lineages that were started from individual *set-2*/+ mothers. We first used this data set to determine the RNA accumulation changes across generations, considering lineages as independent replicates and comparing the later-generation samples (F2, F4 fertiles, and F4 steriles) to the ancestor samples (P0). This confirmed that loss of germline identity observed in *set-2* mutants is associated with significant transcriptional deregulation. Compared to the P0 generation, the number of misregulated genes progressively increased between F2, F4 fertile, and F4 sterile animals, for both up- and downregulated genes (Figure 1B). We observed significantly more upregulated than downregulated genes in all late-generation animals analyzed, as previously reported for healthy *set-2* germlines from animals raised at 20°C (Robert et al., 2014). Therefore, the onset of sterility at 25°C is associated with a progressive deregulation of the germline transcriptome. We also observed that the number of misregulated genes was about 3 times more abundant in sterile than in fertile F4 animals, suggesting that animals remained fertile when deregulation of the transcriptome was moderate, but not when it became more pronounced.

**Figure 1.**
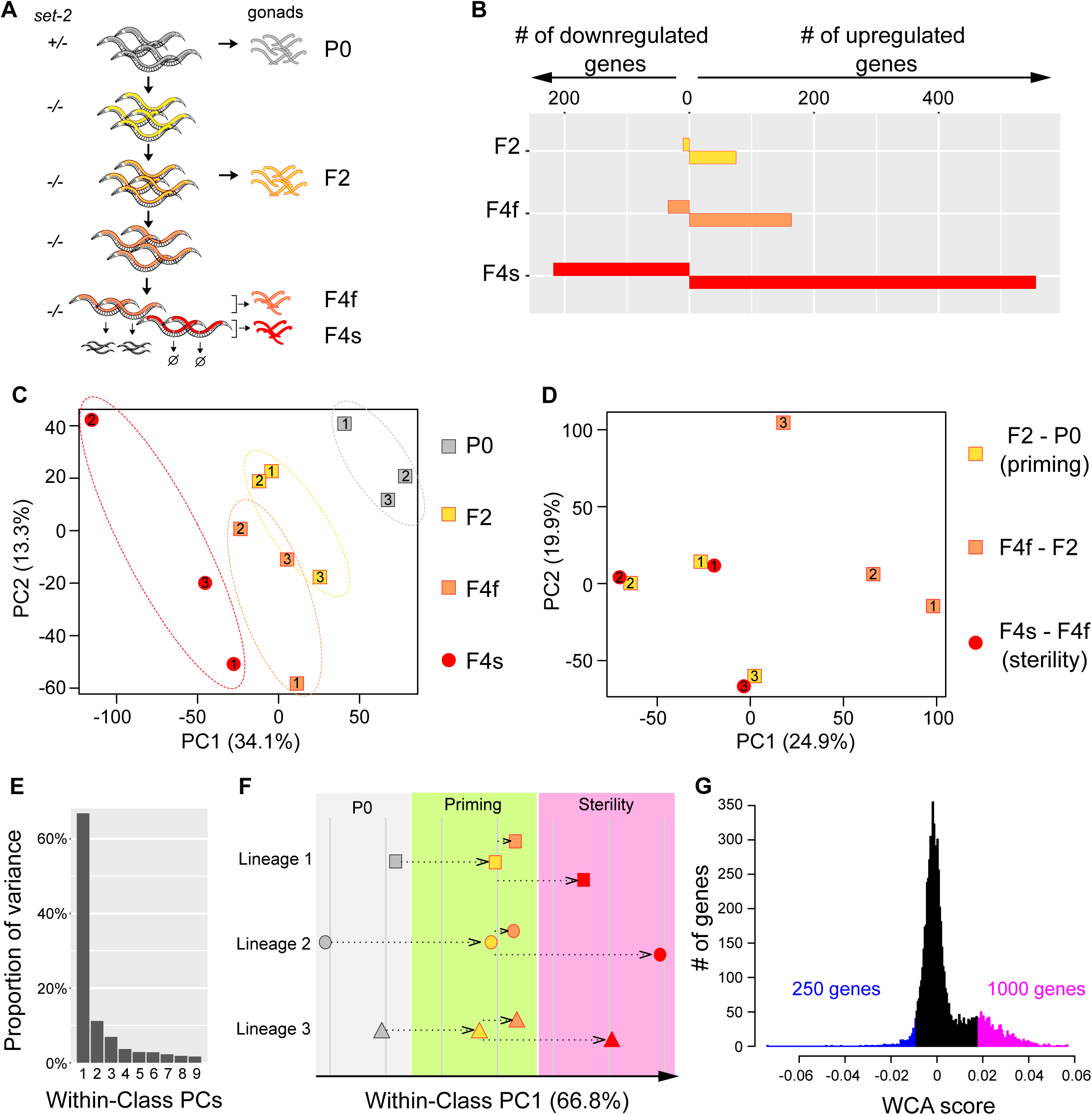
Multivariate analysis of transgenerational transcriptional changes in *set-2* mutant germlines as they progress to sterility. (A) Experimental design of the strategy used to determine transcriptional changes that take place in the gonads of *set-2* mutant animals as they progressively lose their germline identity. Gonads were dissected from P0 *set-2(+)*/*set-2(-)* hermaphrodites, from *set-2(-)*/*set-2(-)* hermaphrodites, and from fertile and sterile F4 *set-2(-)*/*set-2(-)* hermaphrodites for RNA-seq analysis. (B) Number of genes showing differential expression as compared to P0 at a conservative statistical threshold (adjusted *p* < 10^-6^). (C) Principal Component Analysis (PCA) on expression levels. Circles and squares correspond to samples and are numbered according to their lineage. (D) PCA performed on expression differences between samples of the same lineage. Circles and squares correspond to the indicated pairwise differences, numbers indicate lineages. (E-G) Results of Within-Class Analysis (WCA) obtained by considering each lineage as a class. (E) Proportion of within-class variance explained by each component. (F) Coordinates of samples on the first component (one symbol per lineage, same color scheme as in panel C). (G) Histogram showing the distribution of gene coefficients on the first components. Genes having extreme coefficients (colored in blue and magenta) are the major contributors to the orientation of the first component. These coefficients are referred to as “WCA scores” in the main text.

### Multivariate analysis identifies two consecutive steps of transcriptomic deregulation

To visualize the transgenerational progression of gene expression changes in *set-2* mutant germlines, we applied Principal Component Analysis (PCA) on the expression matrix of all genes, and plotted all 12 samples along the first 2 components (Figure 1C). As expected from our initial analysis, the distance relative to the ancestor samples increased over generations and was highest among F4 sterile samples. In addition, and rather unexpectedly, we observed a significant dispersion of the 3 lineages. Inter-lineage distance was more pronounced among F4 fertile samples than among P0 or F2 samples, and was highest among F4 sterile samples (dotted ellipses on Figure 1C). This divergence indicates that transcriptional deregulation did not occurr similarly in the different lineages. Rather, in each lineage, loss of SET-2 resulted in distinct gene expression changes.

To explore this, we computed for each lineage and for all genes the expression differences (log ratios) between F2 and P0 samples, corresponding to early changes; between F4 fertile and F2 samples, corresponding to later changes that did not compromise fertility; and between F4 sterile and F4 fertile samples, corresponding to changes associated with loss of germline function. We applied PCA on the resulting matrix and visualized expression changes along the first 2 principal components (Figure 1D). To our surprise, within each lineage, the early changes (F2-P0) mapped very close to the later changes associated with sterility (F4 sterile-F4 fertile). This was true for all 3 lineages. In contrast, late changes that were not associated with sterility (F4 fertile-F2) mapped further away and were therefore different. This provided a key insight: across the 3 lineages, unique patterns of transcriptomic deregulation occurred early without compromising fertility; in later generations these patterns of deregulation persisted and led to sterility. We therefore call this early step a “priming” process.

### Deregulation of a specific set of genes defines a path to sterility

The analysis above shows that, for each lineage, a set of genes was deregulated in the priming step and further deregulated in a similar direction in sterile F4 animals. We therefore investigated if a common set of genes had experienced this 2-step deregulation towards sterility in all 3 lineages. We first discarded genes that were poorly expressed in all samples, because their variation may simply be due to varying background signal. To determine a meaningful threshold of minimal expression, we plotted the distribution of all genes according to i) their maximal expression level in all samples and ii) their standard deviation of expression across all samples, which could be high due to either meaningless background signal variation or meaningful biological differences between samples. The resulting 2-dimensional density plot revealed 2 subpopulations of genes (Figure S1): a major population that is poorly expressed in all samples, and a secondary population with high expression in at least 1 sample. For the poorly expressed genes, standard deviation increased monotonously with their maximal expression level, as expected for background variation. A different pattern was observed for highly expressed genes (maximal expression greater than 5). First, their standard deviation of expression was not correlated with maximal expression level. This is expected if change in expression is dependent on SET-2 loss and not on background variation. Second, standard deviation was low for the majority of these genes and high for a small subset. This observation is expected if SET-2 acts only on a subset of genes. We therefore chose an arbitrary cutoff of minimal expression that discarded the first subpopulation (red line in Figure S1), leaving a reduced set of 7,238 genes (Table S1).

We next considered each lineage as a class and performed Within-Class Analysis (WCA) on the expression matrix. This method (also called Within-Group Analysis) is a derivative of PCA and is adapted to explore a data set after accounting for the variation due to a known factor (lineage in our case). Although it is particularly powerful to study ecology, its potential for transcriptomic analysis has also been illustrated (Baty et al., 2006). As shown in Figure 1E, the first component of WCA captured most of the Within-Class variation (66.8%) and the subsequent components were not individually informative. We therefore plotted the position of samples along the first component only (Figure 1F). This revealed a continuum of transcriptome evolution that occurred in all 3 lineages, first during the priming step and then towards sterility. This first component of WCA therefore defines a path to sterility. To identify which genes vary along this path, we plotted the distribution of the coefficients that define this component, which we called “WCA scores”. Near-zero scores correspond to genes that do not vary along the path. We examined the transcriptional evolution in each lineage of genes with the largest WCA scores. Out of the 9 genes with the most negative WCA scores, 6 of them were systematically downregulated between P0 and F2 animals and 8 were more strongly downregulated between F2 and sterile F4 than between F2 and fertile F4 (Figure S2). Similarly, out of the 9 genes with the most positive WCA scores, 8 were systematically upregulated between P0 and F2 and all 9 were more strongly upregulated between F2 and sterile F4 than between F2 and fertile F4 (Figure S3). These observations confirm that our WCA analysis identified genes with similar transcriptional trajectories in the 3 lineages. Consistent with the bias towards upregulation in Figure 1B, we observed an asymmetric distribution of WCA scores, with many genes having large positive scores (Figure 1G). We arbitrarily set negative and positive thresholds to define extreme scores. We hereafter refer to the 250 and 1,000 genes with the largest negative and positive scores as “downregulated contributors” and “upregulated contributors”, respectively (Figure 1G and Table S1). These genes make the largest contributions to within-class PC1, and their deregulation is tightly associated with the loss of germline maintenance in all 3 lineages.

### Repression of germline genes and expression of somatic programs are priming events during the loss of germ cell fate in *set-2* mutant germlines

Previous work showed that loss of germ-cell identity in *C. elegans* gonads depleted of PRC2 or P-granule components is associated with decreased expression of germline genes and increased expression of somatic genes (Knutson et al., 2017; Gaydos et al. 2012; Campbell and Updike 2015). To test whether the process of transgenerational loss of fecundity in *set-2* mutants involves similar patterns of gene deregulation, we compared the list of downregulated and upregulated contributors with published lists that categorized genes according to their expression profile in wild-type animals (Knutson et al., 2017). This comparison revealed that downregulation of germline-specific genes and upregulation of soma-specific genes are priming events, as they were detectable before sterility was observed in later generations (Figure 2A and Table S1). Downregulated germline-specific contributors include genes required for meiosis (*apc-10, him-5*), the *wago-4* argonaute-encoding gene, and genes that encode P-granule components (*cey-2, cey-3 and pgl-3).*

**Figure 2:**
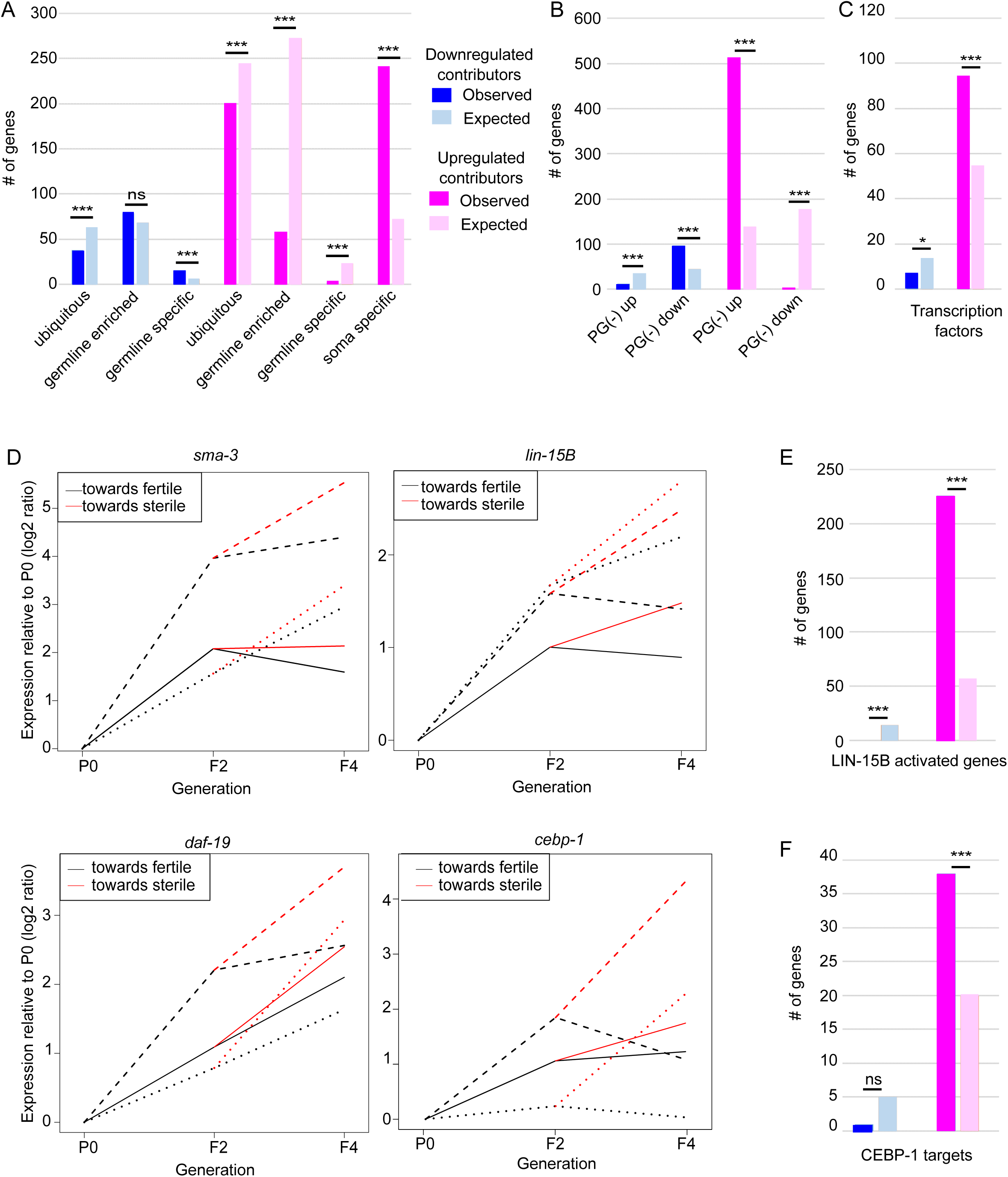
Functional classification of contributors to *set-*2 dependent loss of germline identity. (A) Classification of contributors according to their expression profiles in wild-type animals. Lists of down- and upregulated contributors were compared to lists of ubiquitous, germline-enriched, germline-specific, and soma-specific genes defined in (Knutson et al., 2017). Statistical significance was assessed using a hypergeometric test. Downregulated contributors are depleted for ubiquitous genes (p = 2.3e-4) and enriched for germline-specific genes (p-value = 2e-4); upregulated contributors are depleted for ubiquitous genes (p = 1.3e-4), germline-enriched genes (p = 2.41e-119), and germline-specific genes (2.41e-119), and enriched for soma-specific genes (p= 2.96e-78). * p-value < 0.05, ** p-value < 0.01, *** p-value < 0.001. (B) Comparison of lists of contributors with genes up- or downregulated in the gonads of adult worms in which P granules had been depleted by RNAi for 2 generations (PG(-); Knutson et al., 2017). Color code for down- and upregulated contributors is the same as in (A). Downregulated contributors are depleted for genes upregulated in PG(-) (p = 5.5e-06), and enriched in genes downregulated in PG(-)(p = 1.1e-15); upregulated contributors are depleted for genes downregulated in PG(-) (p-value = 1.4e-90), and enriched in genes upregulated in PG(-) (p-value = 1.4e-218). (C) Comparison of contributors with transcription factors (list wTF3.0). Color code for down- and upregulated contributors is the same as in (A). Statistical significance (hypergeometric test): p-value = 0.03 for under-enrichment of downregulated contributors in transcription factors and p-value = 1.2e-8 for over-enrichment of upregulated contributors in transcription factors. (D) Relative expression to P0 of *sma-3* transcripts, *lin-15B* transcripts, *daf-19* transcripts, and *cebp-1* transcripts in F2, fertile F4 and sterile F4. The 3 independent lineages analyzed are represented by distinctive line formats. For each gene, expression relative to P0 increases more when F4 animals became sterile than when they remain fertile. (E) Comparison of contributors with LIN-15B-activated genes. Color code for contributors is the same as in (A). Statistical significance (hypergeometric test): p-value = 3.4e-7 for under-enrichment of downregulated contributors in LIN-15B-activated genes and p-value = 1.7e-93 for over-enrichment of upregulated contributors in LIN-15B-activated genes. (F) Comparison of contributors with CEBP-1 targets. Color code for contributors is the same as in (A). Statistical significance (hypergeometric test): p-value = 0.2 for under-enrichment of downregulated contributors in CEBP-1 targets and p-value = 4.8e-5 for over-enrichment of upregulated contributors in CEBP-1 targets.

Further analysis revealed that *set-2* inactivation and loss of P-granule components share similar transcriptional signatures (Figure 2B). We observed a significant overlap for both downregulated and upregulated contributors (Table S2; hypergeometric p-value = 1.11e-15 and 1.18e-219, respectively). Detailed investigation of the overlap between P-granule and *set-2* dependent regulation of transcriptional programs showed that downregulated contributors that are also downregulated in P-granule depleted germlines are enriched in both ubiquitous and germline genes (Table S2; hypergeometric p-value = 2.4e-4 and 5.4e-6). By contrast, upregulated contributors also upregulated in the P-granule depleted germlines show no preference for a specific expression pattern (Table S2). For instance, of the 240 soma-specific genes that are upregulated contributors in *set-2* mutant germlines, only 133 are upregulated in P-granule-depleted germlines (Table S2; hypergeometric p-value = 0.07). Therefore, SET-2 and P-granules have both common and unique functions in the repression of somatic gene expression in the germline.

Gene ontology analysis of contributors involved in priming loss of germ-cell identity did not highlight any strongly enriched biological process or pathway. Upregulated soma-specific contributors are involved in a wide variety of biological process including locomotion, nervous system development, axon extension and guidance, morphogenesis, and regulation of pharyngeal pumping. We nonetheless observed that of the total number of transcription factors found in *C. elegans* (wTF3.0; Fuxman Bass et al. 2016), 389 were present in the list of 7,238 genes that we analyzed by WCA, and 101 of these were identified as contributors (1.5-fold enrichment; p = 6e-06) (Figure 2C and Table S3). These include genes encoding SMA-3, an R-SMAD in the TGF-β signaling pathway, LIN-15B, a THAP transcription factor known to antagonize PRC2 activity (Lee, Lu, and Seydoux 2017), DAF-19, an RXF family transcription factor, 7 CEH homeodomain transcription factors, 22 Nuclear Hormone Receptors (NHRs), and CEBP-1, the worm homologue of mammalian CCAAT/enhancer-binding protein (C/EBP). In all 3 lineages, these genes displayed increased expression between P0 and F2 and a stronger increase in expression between F2 and sterile F4 than between F2 and fertile F4 (Figure 2D).

Further analysis suggests that the LIN-15B and CEBP-1 regulatory networks are activated in *set-2* mutant germlines, and that this activation is a priming event in loss of germline identity. Ectopic expression of LIN-15B in primordial germ cells (PGC) results in activation of 452 genes (Lee et al., 2017), of which 226 are present in the list of upregulated contributors (Figure 2E, hypergeometric p-value = 1.71e-93; Table S3). Likewise, in somatic cells CEBP-1 binds 212 protein-coding genes mostly within promoter regions (K. W. Kim et al. 2016), and 38 of these are also present in the list of upregulated contributors in *set-2* mutants (Figure 2E; hypergeometric p-value = 2.5e-28). Interestingly, mammalian C/EBP1 is a key factor in cellular reprogramming (Bussmann et al. 2009; Di Stefano et al. 2014; Sadahira et al. 2012; Xie et al. 2004).

### Progressive upregulation of X-linked genes is a priming event for loss of germline identity in *set-2* mutant gonads

Examination of the genomic distribution of genes identified by WCA revealed that upregulated contributors are strongly enriched on the X chromosome (Figure 3A, hypergeometric p-value = 6e-193), and that these X-linked genes have a significantly higher WCA score than autosomal genes (Figure 3B). Upregulation of X-linked genes is therefore likely to be a priming event driving loss of germline identity. MES-2, MES-3 and MES-6 proteins that form the *C. elegans* version of PRC2 cooperate with MES-4 to repress the X chromosome. Comparison of genes misregulated in the absence of both *mes-2* and *mes-4* (*mes-2;mes-4* double mutants) and our lists identified 188 commonly upregulated genes (Figure 3C; hypergeometric p-value = 2.34e-97, Table S4), mostly located on the X chromosome (137 genes, hypergeometric p-value = 3.2e-21). Conversely, of the 18 commonly downregulated genes, all are found on autosomes (Figure 3D; hypergeometric p-value = 5.84e-14). Common misregulation of target genes is unlikely to reflect regulation of *mes-2* or *mes-4* expression levels by SET-2, since neither of these two genes was identified by WCA. Similarly, *set-2* is not a transcriptional target of PRC2 (Gaydos et al., 2012). Rather, our results suggest that PRC2/MES-4 and SET-2 act in parallel pathways to repress the X chromosome and promote proper expression of a subset of autosomal genes.

**Figure 3:**
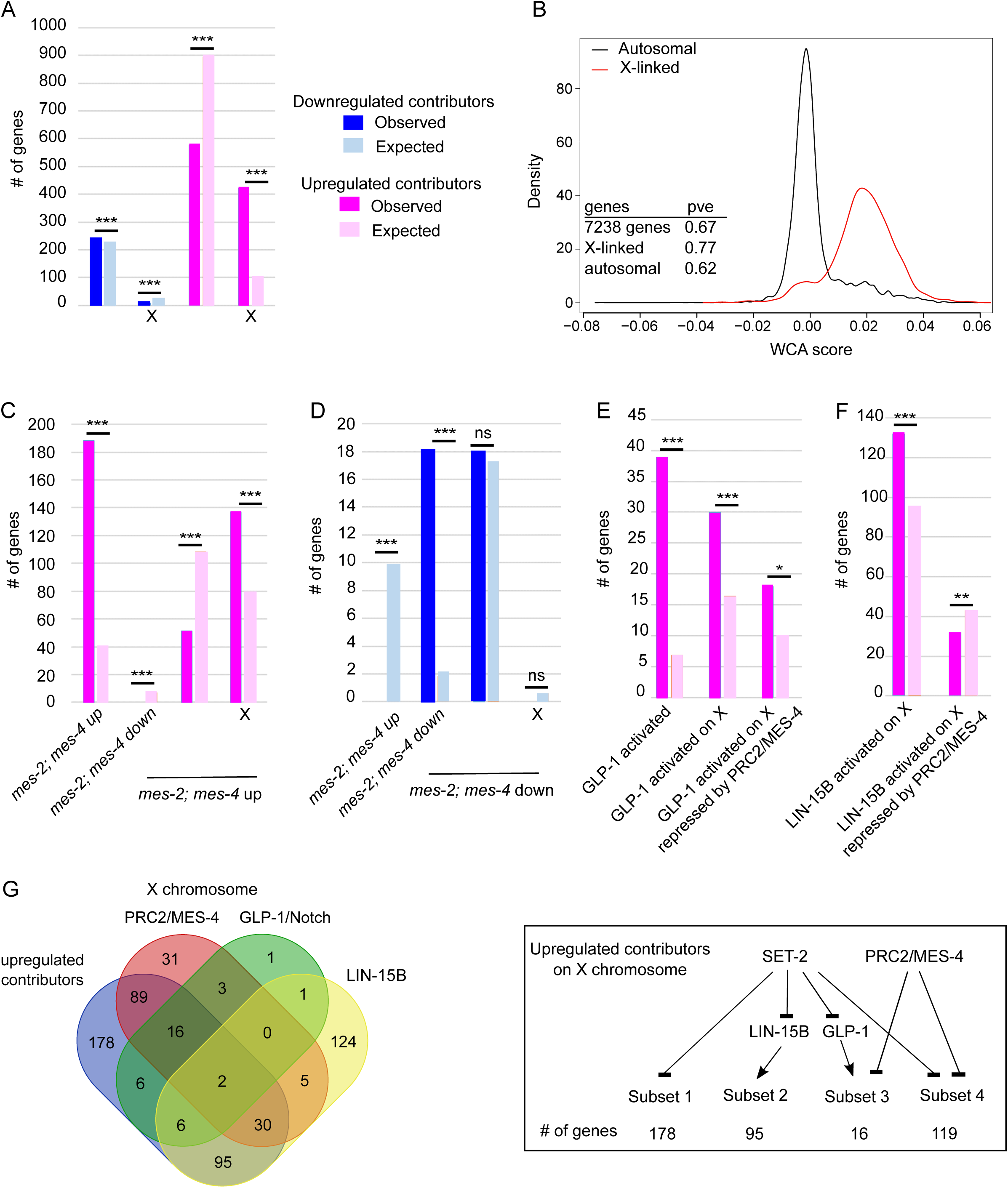
Chromosome X desilencing is a priming event for loss of germline identity. (A) Chromosome distribution of contributors. Statistical significance (hypergeometric test): p = 4.4e-5 and 6.1e-193 for distribution of down- and upregulated contributors, respectively. * p-value < 0.05, ** p-value < 0.01, *** p-value < 0.001. (B) Distribution of WCA score according to location of the genes on autosomes or the X chromosome. (pve = percent of variance explained). (C) Comparison of upregulated contributors with *mes-2*; *mes-4* up- and downregulated genes and the chromosome distribution of the upregulated class. Color code is the same as in (A). Statistical significance (hypergeometric test): p = 2.34e-97 for over-enrichment in *mes-2*; *mes-4* upregulated genes, p = 3.6e-4 for under-enrichment in *mes-2*; *mes-4* downregulated genes, p-value = 3.2e-21 for chromosome distribution. (D) Comparison of downregulated contributors with *mes-2*; *mes-4* up- and down-regulated genes and the chromosome distribution of the downregulated class. Color code is the same as in (A). Statistical significance (hypergeometric test): p-value = 3.9e-5 for under-enrichment in *mes-2*; *mes-4* upregulated genes, p-value = 5.84e-14 for over-enrichment in *mes-2*; *mes-4* downregulated genes, p-value = 0.5 for chromosome distribution. (E) Comparison of upregulated contributors with genes activated by GLP-1. Color code is the same as in (A). Statistical significance (hypergeometric test): p-value = 1.25e-24 for over-enrichment in GLP-1 activated genes, p-value = 7e-6 for over-enrichment in X-linked GLP-1 activated genes, p-value = 1e-2 for over-enrichment in X-linked GLP-1 activated genes repressed by PRC2/MES-4. (F) Comparison of upregulated contributors with genes activated by LIN-15B. Color code is the same as in (A). Statistical significance (hypergeometric test): p-value = 7.7e-9 for over-enrichment in X-linked LIN-15B-activated genes, p-value = 8e-3 for over-enrichment in X-linked LIN-15B-activated genes repressed by PRC2/MES-4. (G) Comparison of X-linked upregulated contributors with genes repressed by PRC2/MES-4 and genes activated by GLP-1/NOTCH or LIN-15B. Summary of connections between the regulatory networks that regulate various classes of X-linked genes in *set-2* mutant germlines.

### Ectopic expression of GLP-1/Notch signaling and the LIN-15B transcription factor contributes to derepression of X-linked genes in *set-2* mutant germlines

PRC2-dependent repression of the X chromosomes in the germline is counteracted by increased GLP-1/Notch signaling: in worms bearing the gain-of-function allele *glp-1(ar202)*, increased signaling induces expression of specific genes normally repressed by PRC2 (Seelk et al., 2016). We found that *glp-1* is over-expressed in all 3 lineages across generations (WCA score = 0.0054), and GLP-1/Notch targets are upregulated contributors in our analysis (39 genes representing a 5.65-fold enrichment; hypergeometric p-value = 1.25e-24, Figure 3E). Of these targets, 30 are located on the X chromosome (hypergeometric p-value = 7.5e-06, Table S1), and 18 of these are upregulated in germlines lacking both MES-2 and MES-4 (hypergeometric p-value = 1e-3). Altogether, these data support a model in which SET-2 represses a subset of X-linked genes through inhibition of GLP-1/Notch, and some of these same genes are also PRC2/MES-4 targets (Figure 3G).

An additional subset of upregulated contributors on the X chromosome is enriched in targets of the LIN-15B transcription factor (Figure 3F; hypergeometric p-value = 7.71e-9). This same set of genes is slightly depleted for PRC2/MES-4 targets (hypergeometric p-value = 7e-3), suggesting that SET-2 represses their expression through inhibition of LIN-15B and independently of PRC2.

### The onset of sterility can be delayed by countering the upregulation of individual contributors

Our analysis identified a set of genes whose deregulation occurs in early generations of *set-2* mutant germlines without compromising fertility, but whose further deregulation in later generations leads to sterility. Focusing on upregulated contributors, we reasoned that if increased expression of these genes is an essential priming step in the process leading to sterility, then their downregulation before animals have become sterile might delay the onset of sterility. To test this, we used RNAi to knock-down candidate genes from our WCA list in *set-2* mutant animals and scored the onset of sterility by counting the number of progeny laid over 3 days compared to *set-2* mutant animals not treated with RNAi (Figure 4A). We observed that in control experiments, growth on the HT115 *E. coli* strain used for RNAi feeding delays the onset of sterility compared to growth on the OP50 strain routinely used for culturing, consistent with bacterial diet influencing fertility and other phenotypes (Watson, Yilmaz, and Walhout 2015; Heestand et al. 2018). We first tested 29 upregulated contributors. RNAi knock-down of 9 of these significantly and reproducibly delayed the onset of sterility in *set-2* mutant animals over generations (Figures 4B, 4C, Table S5). These include *(i) cebp-1, daf-19, attf-5*, and *somi-1*,encoding transcription factors; (ii) *pek-1*, encoding a homologue of mammalian EIF2A kinase; (iii) *avr-14*, encoding a glutamate-gated chloride channel; (iv) *crb-1*, involved in epithelial polarization; (v) the *aldo-1* aldolase gene; and (vi) the *fhod-1* formin gene. While the number of progeny counted on *set-2* control plates (no RNAi) dramatically decreased between the F4 and F8 generation, and animals became completely sterile at the F14 generation, 16 to 66% (depending on the targeted gene) of the RNAi plates still contain fertile animals in the F14 generation and beyond. As previously reported for mutants showing progressive loss of sterility, loss of fecundity and brood sizes were extremely variable (Robert et al., 2014;Yanowitz, 2008). RNAi knock-down of 11 additional genes (*his-8, ceh-43, hbl-1, lin-59, nhr-57, sop-2, sor-1, ckk-1, eef-1A.2, glp-1*, and *rpl-11.2)* resulted in early onset of sterility that prevented further investigation of their role (data not shown). Finally, for the remaining 9 genes (*flh-2, gfi-3, nhr-48, jmjd-3.1, utx-1, kgb-1, puf-9, miz-1*, and *ncam-1)*, no significant transgenerational effect on fertility was observed after RNAi treatment. For these genes, we cannot rule out that the absence of an effect is due to failure of the RNAi treatment to knock-down their expression.

**Figure 4:**
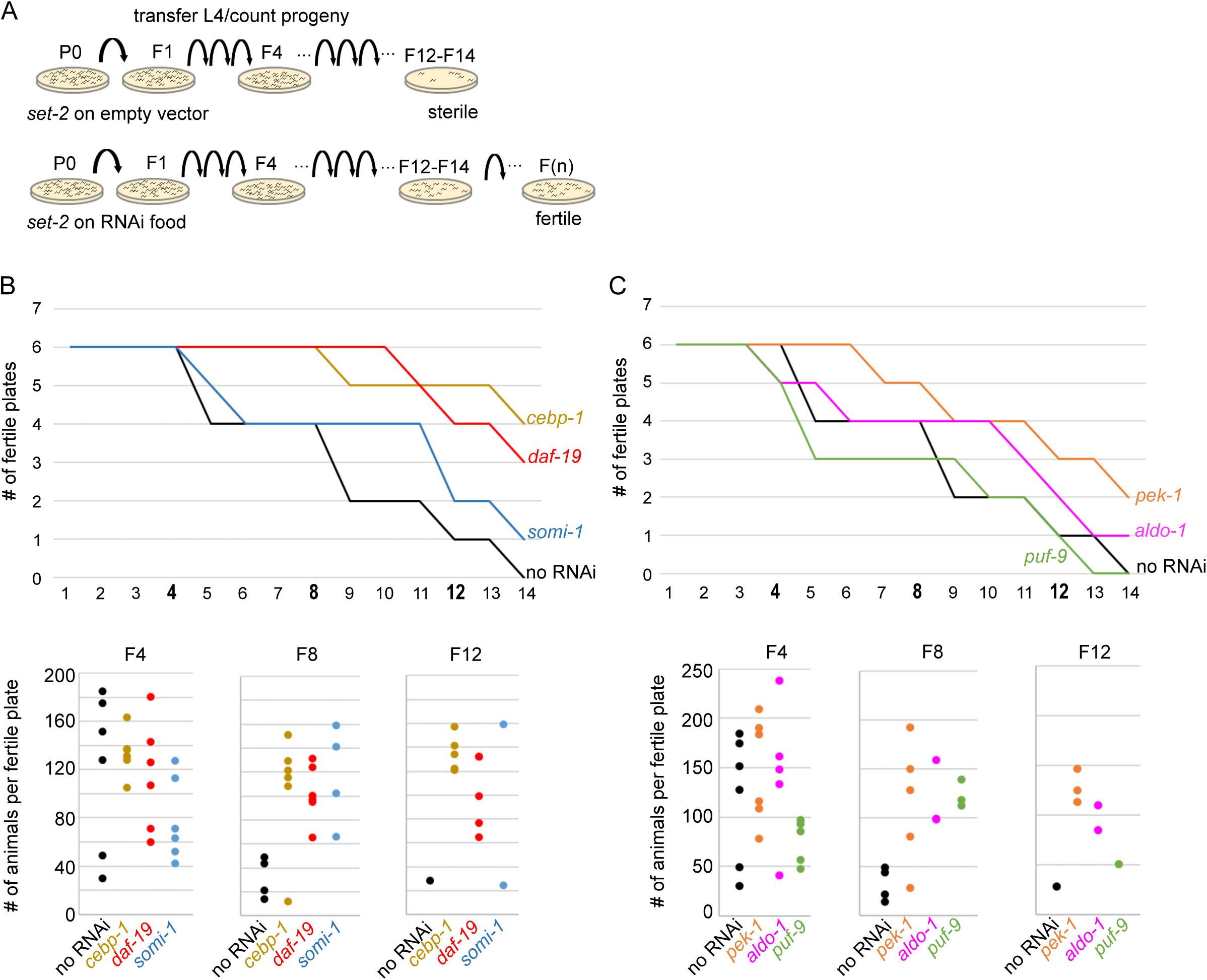
Onset of sterility is delayed by RNAi knock-down of upregulated contributors. (A) Experimental design of the RNAi strategy to test the effect of knock-down of upregulated contributors in *set-2* dependent loss of germline identity. (B) *cebp-1, daf-19*, and *somi-1* RNAi knock-down delays the onset of sterility compared to animals fed with an empty RNAi vector (no RNAi). Number of fertile plates (with less than 20 animals) per generation (upper panel) and number of animals per fertile plate in F4, F8, and F12 (lower panel) are shown. (C) *pek-1* and *aldo-1* but not *puf-9* RNAi knock-down result in a delay in the onset of sterility compared to animals fed with an empty RNAi vector (no RNAi).

### Expression of TGF-β pathway components in *set-2* mutant germlines primes the process leading to sterility

Our identification of the *C. elegans* SMAD SMA-3 as a positive contributor to loss of germ-cell fate raised the possibility that activation of TGF-β signaling in *set-2* mutant germlines could be a priming event. In *C. elegans*, 2 canonical TGF-β signaling pathways have been described, defined by the DAF-7 and DBL-1 ligands that function through distinct serine–threonine kinase receptors and their pathway-specific SMADs (Figure S4A; Savage-Dunn and Padgett 2017). Analysis of the WCA list revealed the presence of components of both pathways, including the Type I receptor DAF-1, R-SMADs (SMA-3, DAF-8 and DAF-14), Co-SMADs (SMA-4 and DAF-3), and downstream transcription factors (SMA-9 and DAF-12). If genes with the top 1500 (instead of 1000) WCA scores are included, this list can be extended to the Type II receptor DAF-4, the extracellular regulator SMA-10, and the transcription factor DAF-5. Upregulated contributors also include KIN-29, a serine/threonine kinase that interacts with the DBL-1 pathway (Maduzia et al., 2005) and OBR-3, which may participate in TGF-β signaling by binding the BMP receptor-associated BRA-1 and BRA-2 proteins (Sugawara et al., 2001). For all these TGF-β signaling components and regulators, transcript levels increased in all 3 lineages between P0 and F2. Furthermore, for at least 2 out of the 3 lineages, expression increased more between F2 and sterile F4 animals than between F2 and fertile F4 animals (Figure 2D, Figure 5A and S4B). RNAi knock-down of *sma-3, sma-9, daf-5, sma-10, kin-29* and *obr-3* in *set-2* mutants resulted in a reproducible delay in the onset of sterility (Figure 5B). Altogether, these results suggest that germline expression of TGF-β pathway components may render the germline responsive to signaling cues from the soma, where the DAF-7 and DBL-1 ligands are expressed.

**Figure 5:**
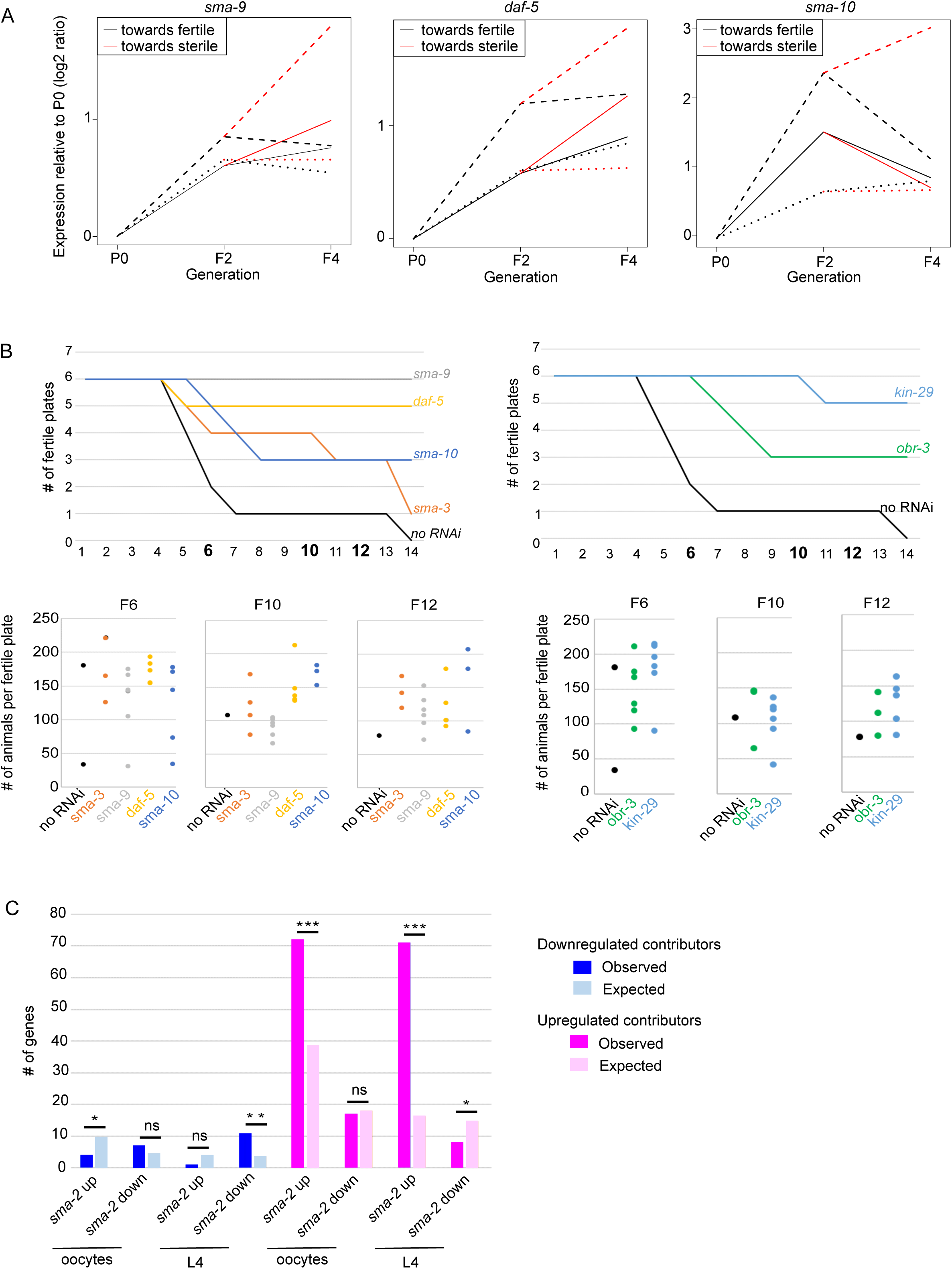
Ectopic TGF-beta signaling may contribute to loss of germline identity. (A) Expression relative to P0 of *sma-9, daf-5*, and *sma-10* transcripts in F2, F4 fertile, and F4 sterile. The 3 independent lineages analyzed are represented by distinctive line formats. (B) *sma-9, daf-5, sma-10*, and *sma-3* RNAi knock-down (left panel) and *kin-29* and *obr-3* RNAi knock-down (right panel) delay the onset of sterility of *set-2* mutant animals compared to animals fed with an empty RNAi vector (no RNAi). Number of fertile plates per generation (upper panel) and number of animals per fertile plate in F6, F10, and F12 (lower panel) are shown. (C) Comparison of contributors with genes up- and downregulated in *sma-2* mutant animals. Statistical significance (hypergeometric test): for downregulated contributors, p-value = 0.03 for under-enrichment in *sma-2* oocyte upregulated genes, p-value = 0.17 for over-enrichment in *sma-2* oocyte downregulated genes, p-value = 0.08 for under-enrichment in *sma-2* L4 upregulated genes, p-value = 0.001 for over-enrichment in *sma-2* L4 downregulated genes; for upregulated contributors: p-value = 5e-8 for over-enrichment in *sma-2* oocyte upregulated genes, p-value = 0.43 for under-enrichment in *sma-2* oocyte downregulated genes, p-value = 1.6e-32 for over-enrichment in *sma-2* L4 upregulated genes, p-value = 0.03 for under-enrichment in *sma-2* L4 downregulated genes. * p-value < 0.05, ** p-value < 0.01, *** p-value < 0.001.

In wild-type animals, TGF-β signaling from neurons regulates both somatic developmental processes in the soma and reproductive processes in oocytes, through temporally and transcriptionally distinct programs. If TGF-β signaling is active in the germline of *set-2* mutant animals, we reasoned that germline misexpression of its target genes should also be observed. Comparison of upregulated and downregulated contributors with lists of TGF-β transcriptional targets (Luo et al. 2010) revealed that upregulated contributors are enriched in genes upregulated in oocytes of *sma-2* mutant animals, as well as a largely non-overlapping set of genes upregulated in *sma-2* mutant L4 animals, prior to oocyte development. By contrast, no enrichment for TGF-β downregulated genes was found (Figure 5C). Analysis of the 2 sets of genes showed that a total of 137 upregulated contributors are repressed by TGF-β signaling either in L4 (71 genes, hypergeometric p-value =1.6e-32) or in oocytes (72 genes; hypergeometric p-value = 5.00e-08; Figure S4C). This was an unexpected finding, since these are genes normally repressed by TGF-β signaling, which is activated in our mutant germlines. These upregulated contributors are enriched in soma-specific genes and LIN-15B activated genes (Figure S4D), suggesting that LIN-15B and TGF-β cooperate at a subset of loci to promote gene expression during loss of germ cell fate. Downregulated players showed no enrichment for TGF-β targets. Altogether, these results suggest a complex network whereby ectopic activation of TGF-β pathway components in the germline of *set-2* mutant animals results in transcriptional activation of genes that are normally repressed by TGF-β signaling in L4, and to a lesser extent in oocytes.

## Discussion

In this study we used transgenerational expression profiling to identify genes that prime the loss of germ cell identity in the absence of the SET-2/SET1 H3K4 methyltransferase. Our analysis shows that further misregulation of this same set of genes is responsible for sterility in later generations, thereby identifying contributors in a two-step deregulation process leading to sterility in independent lineages.

Our work illustrates the complementarity between multivariate analysis, which was key to identifying these contributors, and the pairwise comparisons commonly used in transciptomics. In a pairwise comparison, for each gene one considers the null hypothesis that its expression level is unaffected by the perturbation under study (absence of SET-2 in our case). This approach efficiently identifies target genes as those displaying fold-changes and statistical significance above a chosen threshold value (as in Figure 1B). Each gene therefore corresponds to a pronounced quantitative effect resulting from the experimental perturbation. However, because the null hypothesis cannot always be reformulated adequately, this approach is limited when additional factors contribute to variation, for example in this study the lineage from which the animals were derived. Within-class analysis has the advantage of exploiting the variation of thousands of genes simultaneously, while also accounting for these additional variables. Using WCA we were able to extract a signature of global variation and identify genes contributing to this signature. Even if the identified genes varied modestly in terms of fold-change, their variation was highly correlated with the gradual progression towards sterility. As a validation of WCA to identify genes involved in maintenance of fertility, it is noteworthy that out of 24 positive contributors that we could test using RNAi, 15 were shown to delay onset of fertility when inactivated.

Previous studies showed that loss of PRC2/MES-4 and P-granule components results in a global deregulation of the germline transcriptome (Knutson et al. 2017; Updike et al. 2014; Gaydos et al. 2012). However, these studies could not distinguish between genes whose altered expression is a priming event and genes whose altered expression is a consequence of loss of germline identity. Here we show that priming involves concurrent upregulation of somatic genes and downregulation of germline genes, and that upregulated genes are greatly enriched on the X chromosome. The role of PRC2/MES-4 in repressing the X chromosome in the germline is well characterized (Bender et al. 2006; Gaydos et al. 2012; Rechtsteiner et al. 2010). We found that affected genes on the X fall into two classes: a large class consisting of genes repressed by SET-2 independently of PRC2, and a smaller class repressed by SET-2 and PRC2/MES-4. The second class is consistent with a common regulatory role for SET-2 and PRC2/MES-4 on the X chromosome and provides an explanation for genetic data showing that depletion of SET-2 enhances the sterility of *mes* mutants (Xu and Strome 2001).

While SET-2 is responsible for deposition of H3K4me3 in the germline (Li and Kelly 2011; Xiao et al. 2011), immunostaining experiments showed that marks of active chromatin including H3K4me3 are absent from the X chromosomes in proliferating and early meiotic germ cells, and only appear on the Xs during oogenesis (Kelly et al. 2002). Therefore, loss of H3K4me3 from the X chromosome of *set-2* mutants is unlikely to play a causal role in loss of repression on this chromosome. Rather, loss of H3K4me3 from autosomes may indirectly affect chromatin structure on the X. Consistent with such a model, we previously observed global changes in H3K9me3 and H3K27me3 in *set-2* mutant germlines (Robert et al., 2014). Regardless of the exact distribution of these marks on the X in *set-2* mutant germlines, which remains to be established, these results emphasize the importance of maintaining global repression of the X in order to preserve germ cell identity.

We observed significant overlap between genes misregulated following loss of P-granule components, and contributors to the onset of sterility in *set-2* mutants. This and the observation that *set-2* mutant germlines nearing sterility lose P-granules (Robert et al, 2014) suggest that *set-2* contributes to the maintenance of germline identity by stabilizing P-granules. However, while P granules are necessary to downregulate spermatogenesis genes in adult germ cells (Knutson et al., 2017), spermatogenesis genes are not affected by loss of SET-2. This suggests that additional mechanisms participate in the loss of germ cell identity in P-granule-defective germlines, or that misregulation of spermatogenesis genes in the absence of P granules may not be directly responsible for the loss of germline identity. We note that almost half of the upregulated contributors in *set-2* dependent loss of germline identity, including the majority of TGF-β pathway components, are not misregulated in the absence of P granules, further supporting that the role of SET-2 in maintaining germline identity is not restricted to stabilization of P granules.

The observation that transcription factors are significantly over-represented in the set of upregulated contributors is consistent with the central role of transcription factor networks in reconfiguring cellular identity. One of the contributors we identified is CEBP-1, which is required for adult sensory axon regeneration and neuronal stress responses (Bounoutas et al. 2011; Yan et al. 2009; K. W. Kim et al. 2016). Importantly, known targets of CEBP-1 were also identified as significant contributors in the loss of cell identity. In mice, the C/EBP family of transcription factors regulates cell proliferation and differentiation (Nerlov 2007), and C/EBPα enhances reprogramming of B cells (Bussmann et al., 2009; Sadahira et al., 2012; Xie et al., 2004) at least in part by increasing chromatin accessibility to reprogramming factors (Di Stefano et al., 2014). Its expression in the germline could therefore prime cells to respond to additional transcription factors, leading to neuronal differentiation.

We also identified numerous TGF-β pathway components, although the ligands DAF-7 and DBL-1 were not present in our list of contributors. Downstream targets of TGF-β signaling in somatic tissues were also identified as upregulated contributors, suggesting that the pathway is active despite the absence of expression of known ligands in the germline. Signaling from ligands expressed in somatic tissues to germline responsive cells could provide cues that initiate somatic development. Alternatively, signalling in mutant germlines may be initiated independently of ligand binding (Massagué 2012). Unexpectedly however, TGF-β target genes contributing to priming loss of germ cell identity in the absence of *set-2* were repressed, rather than activated in *set-2* deficient germlines. Several factors may account for this apparent paradox. First, because SMADs have weak binding affinity for DNA (Massagué 2012), their transcriptional role depends on robust interaction with other transcription factors and chromatin-associated proteins, resulting in cell-type and context-dependent cellular responses to TGF-β activation (Derynck and Zhang 2003; Ikushima and Miyazono 2010; Massagué, Seoane, and Wotton 2005) that may be substantially different between normal, healthy somatic tissues and mutant germlines. Second, alternative signaling mechanisms other than the canonical Sma/Mab TGF-β family signaling pathways could be activated in the context of *set-2* mutant germlines undergoing cell conversion (Savage-Dunn and Padgett, 2017).

Our analysis was carried out on a population of germ cells. Therefore, we do not know whether the progressive transcriptional changes we observed reflect increased expression over time of individual genes in a few cells, or across the entire population. However, we observed that a neuronal marker is not expressed in all cells of *set-2* mutant gonads (Robert et al. 2014). Therefore, we favor a model in which the absence of SET-2 results in an increased tendancy for somatic genes to be expressed in individual germ cells, and the percentage of germ cells that misexpress somatic genes increases over generations, eventually compromising fertility (Losick and Desplan 2008).

Decreasing expression of positive contributors to loss of germ cell identity, including *cebp-1* and TGF-β pathway components, was sufficient to delay the onset of sterility, suggesting that these genes individually contribute to the process leading to sterility. Therefore, loss of germ cell fate is driven by misexpression of many different genes whose expression is repressed in wild-type germlines by SET-2.

We observed global changes in chromatin organization in *set-2* mutant fertile germlines, suggesting that these modifications precede the transcriptional changes leading to sterility (Robert et al. 2014). This is consistent with previous observations that histone modifications maintain progenitor cell identity as well as prime them for differentiation, and that extensive alterations of genome-wide chromatin organization are generally much more widespread than initial changes in gene expression during the early phases of loss of cell identity (Koche et al. 2011). In the future it will be informative to correlate the transcriptional reprogramming described here to changes in gene expression in individual germ cells.

## Materials and methods

### Strains and maintenance

Nematode strain maintenance was described previously (Brenner 1974). The strain *set-2*(*bn129*) III/*qC1*(*qIs26*) used in this study is described in (Herbette et al. 2017).

### Transcriptomic analysis

Gonads from *set-2* homozygous worms were collected from 3 independent lineages derived from *set-2*(*bn129*)/*qC1*(*qIs26*) (P0 *set-2*(+)/*set-2*(-)) animals by initially picking 25 *set-2*(+)/*set-2*(-) L4 worms and then cloning single worms from subsequent generations to individual plates. Fertility of each worm was assessed before dissection by monitoring egg laying for 24 hours after young adult stage. Worms that laid any live progeny were considered as fertile. For the F4 sterile samples, only worms lacking visible embryos in the uterus were collected. For each replicate, seventy five to one-hundred gonad arms were dissected from *set-2*(*bn129*)/*qC1*(*qIs26*) (P0 *set-2*(+)/*set-2*(-)) and *set-2*(*bn129*)/*set-2*(*bn129*) (F2 and F4 *set-2*(-)/*set-2*(-)) animals. Dissected gonads were cut at the gonad bend with 30 1/2-gauge needles in egg buffer (pH 7.3, 27.5 mM HEPES, 130 mM NaCl, 2.2 mM MgCl_2_, 2.2 mM CaCl_2_, and 0.528 mM KCl) containing 0.5% Tween 20 and 1 mM levamisole and collected into Trizol. Total RNA was extracted and ribosomal RNA was depleted using an NEBNext rRNA Depletion Kit (Human/Mouse/Rat) (catalog number E6310). Libraries were constructed using an NEBNext Ultra RNA Library Prep Kit for Illumina sequencing (catalog number E7530) and sequenced at the Vincent J. Coates Genomics Sequencing Laboratory at the University of California, Berkeley, using Illumina HiSeq 2500 and 4000 platforms. For differential expression analysis, raw sequences were first mapped to transcriptome version WS220 using TopHat2 (D. Kim et al. 2013). Only reads with one unique mapping were allowed, otherwise default arguments were used. Reads mapping to ribosomal RNAs were then removed. HTSeq (Anders, Pyl, and Huber 2015) was used to build a count table of expression levels per transcript. DESeq2 (Love, Huber, and Anders 2014) was used to normalize read counts across samples, leading to matrix M that contained the normalized expression levels of 20261 genes. DESeq2 was also used to determine genes for which expression at a downstream generation differed from expression at P0, with no consideration for lineage-dependent relations (Figure 1C). *P*-values were adjusted for multiple testing using the Benjamini–Hochberg method as implemented in DESeq2.

### Multivariate analysis

We did all further multivariate analysis using R (www.r-project.org) version 3.4.4. To perform PCA (Figure 1C), we first transformed M by applying × = Log_2_(x + 1) to all of its values, obtaining matrix LM, which we transposed and processed with the prcomp() function. To perform PCA on expression differences (Figure 1D), we constructed a 20261 × 9 matrix (DLM), where rows were genes and columns were expression differences of interest (*e.g*. F2-P0 lineage 1) which we computed by subtracting one column of LM (*e.g.* F2 lineage 1) from another (*e.g.* P0 lineage 1). We then transposed DLM and processed it with the prcomp() function. Before applying WCA, we subsetted expression matrix LM by keeping only the 7238 genes having at least one value greater than 5 (Figure S1). We then processed the resulting matrix with the dudi.pca() function of the ade4 package (Thioulouse et al. 1997) (version 1.7-6, http://pbil.univ-lyon1.fr/ADE-4/home.php?lang=eng;) using parameters scan = FALSE, scale = FALSE and nf = 4. We then processed the resulting object with the ade4::wca() function using lineages as classes and parameters scan = FALSE, nf = 2. The first column of attribute c1 of the resulting object corresponded to the WCA scores reported in text and figures.

### RNAi screen

Bacterial clones containing RNAi feeding vectors were collected in the *C. elegans* RNAi collection (made by J. Ahringer, Source Bioscience). The molecular sequence of insert present in each RNAi clones was checked by sequencing (primer ggtcgacggtatcgataagc) after PCR amplification (single primer in duplicated T7 promoter taatacgactcactataggg) performed directly on colonies. For feeding, bacterial clones were amplified 18 hours at 37°C in LB complemented with Ampicilline (50μg/ml). Transcription (from duplicated T7 promoters) was induced by adding IPTG (1mM final) and growing liquid cultures for 2 additional hours at 37°C. 200 μl of induced cultures were plated on NGM plates complemented with IPTG (1mM). At each generation, 6 L4 animals were transferred on RNAi plates and grown at 25°C for 3-4 days. Progeny was briefly counted every two generations to evaluate animals fertility. Animals were considered as sterile when less than 20 progeny were present on a plate.

## Acknowledgements

We thank Dieu Huong Hoang, Sonia Grzeskowiak and Chloé Exbrayat-Héritier for help with experiments, Laurent Modolo (Pôle Bioinformatique, LBMC) for suggesting multivariate class analysis, developers of R, ade4 package and Ubuntu for their software.

S.S. is supported by NIH grant R01 GM34059, F.P. by ANR grant N° 15-CE12-0018-01, the Fondation ARC and the Centre National de la Recherche Scientifique.

## Author contribution

Conceptualization: Valérie Robert, Gaël Yvert, Susan Strome, Francesca Palladino.

Formal Analysis: Andreas Rechtsteiner and Gaël Yvert.

Funding Acquisition: Susan Strome and Francesca Palladino.

Investigations: Valérie Robert and Andrew Knutson.

Methodology: Valérie.Robert., Gaël Yvert, Susan Strome and Francesca Palladino.

Writing - Original draft preparation: Valérie Robert, Gaël Yvert and Francesca Palladino.

---

**Table S1:** list of 7238 genes, downregulated and upregulated contributors with ubiquitous/germline/soma classification and comparison with genes deregulated following P-granule knock-down. Description of germline-specific downregulated contributors. Description of soma-specific upregulated contributors.

**Table S2:** raw data of hypergeometric test performed on data presented on Figure 2, Figure 3, Figure 5 and Figure S4.

**Table S3**: Comparison with transcription factors (wTF3.0), LIN-15B activated genes and CEBP-1 targets. Description of transcription factors found in upregulated and downregulated contributors.

**Table S4:** Comparison with genes misregulated in mes-2; mes-4 mutants and with genes activated by GLP-1/Notch and LIN-15.B

**Table S5:** RNAi targeting of positive contributors to the WCA PC1 in a *set-2*(*lf*) background. Full list of tested genes.

**Table S6:** comparison with TGF-β targets

**Figure S1: Two-dimensional density plot of 20**,**261 genes according to their maximal expression level (X-axis) and standard deviation (Y-axis) across all samples of the dataset.** Each circle represents a gene. Gene density is represented as heat maps. The red vertical line delimits the set of 7,238 genes showing an expression value greater than 5 in at least one sample and analysed by WCA.

**Figure S2: Expression levels relative to P0 in F2, F4 fertile and F4 sterile for the top 9 most downregulated contributors.** The 3 independent lineages analyzed are represented by distinct line formats.

**Figure S3: Relative expression to P0 of transcripts in F2, fertile F4 and sterile F4 of transcripts of the top 9 most upregulated contributors.** The 3 independent lineages analyzed are represented by distinct line formats.

**Figure S4. TGF-β components upregulation contributes to loss of germline identity in *set-2* mutant animals.** (A) Scheme of the TGF-β pathway present in *C. elegans*. Components upregulated during loss of germline identity are in red. (B) Expression relative to P0 of *daf-4, daf-14, daf-8, sma-4, daf-3, obr-3* and *kin-29* transcripts in F2, F4 fertile and F4 sterile. (C) Overlap between upregulated contributors and genes upregulated in *sma-2* oocytes or *sma-2* L4 stage animals. (D) Comparison of the list of 137 genes upregulated in *sma-2* oocytes or *sma-2* L4 stage animals and *set-2* mutant germlines with various classes of genes. Soma specific genes (p-value = 2e-4) and LIN-15B activated genes (p-value = 5e-4) are significantly over-represented in this dataset. * p-value < 0.05, ** p-value < 0.01, *** p-value < 0.001.

