## Supplementary figures and images for "The *C. elegans* SET-2 histone methyltransferase maintains germline fate by preventing progressive transcriptomic deregulation across generations"

### Supplemental Figure 1

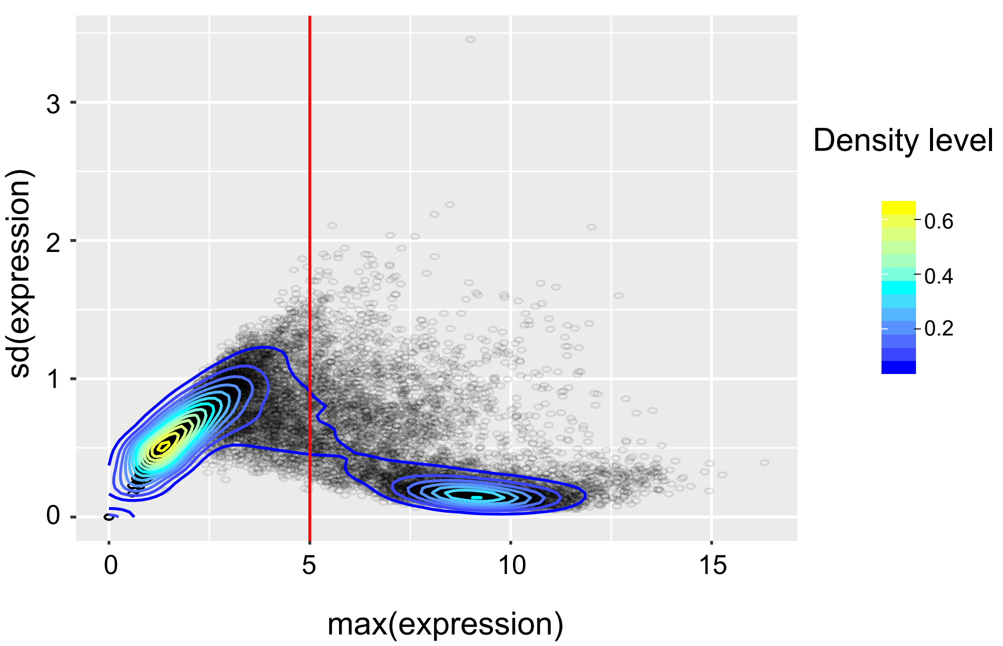

### Supplemental Figure 2

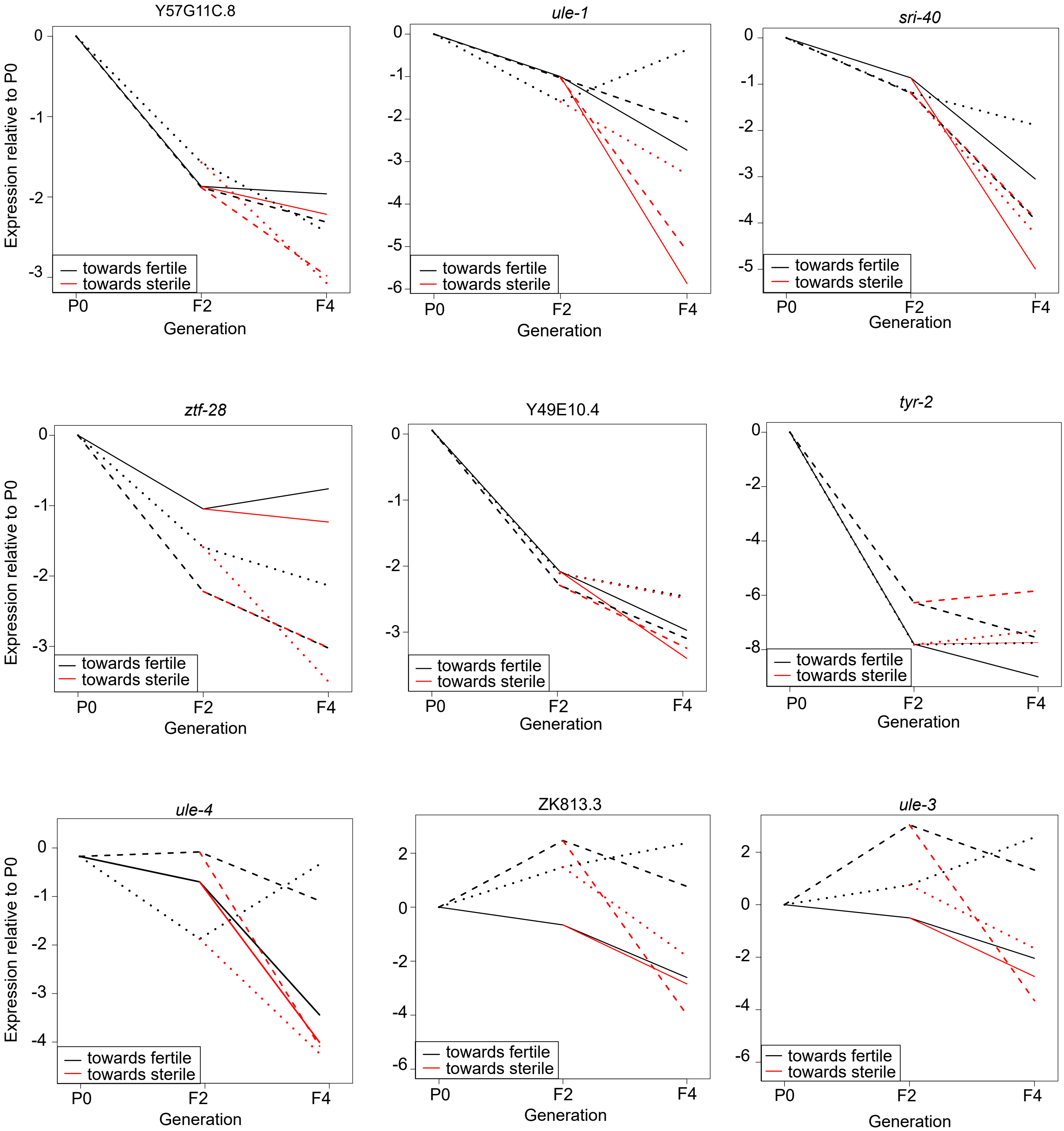

### Supplemental Figure 3

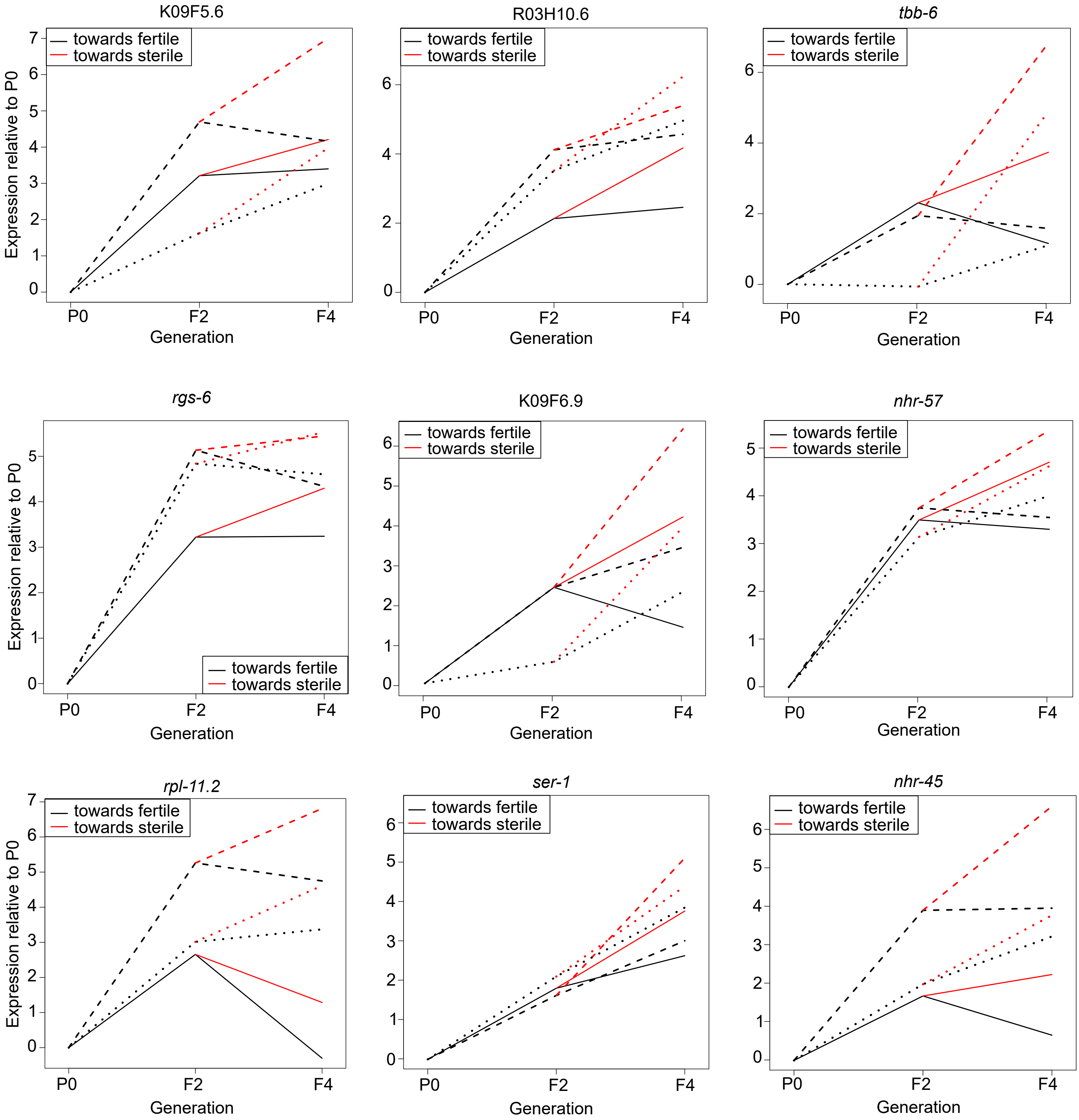

### Supplemental Figure 4

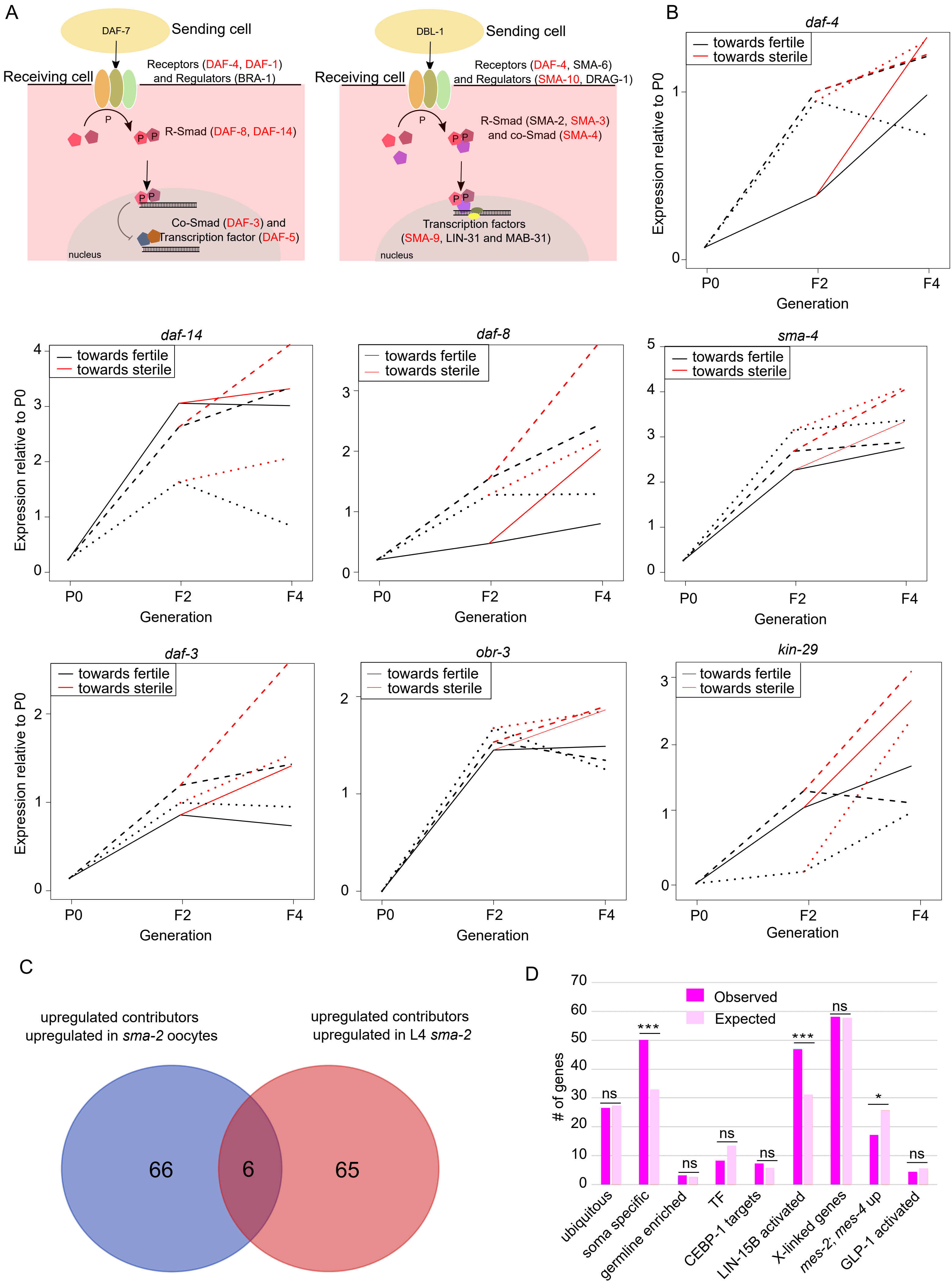
