## Supplemental Table 5 for "The *C. elegans* SET-2 histone methyltransferase maintains germline fate by preventing progressive transcriptomic deregulation across generations"

| Functional class | chr | Generations | | |
| --- | --- | --- | --- | --- |
|  |  | F4 | F8 | F12 |
| Transcription |  |  |  |  |
| *cebp-1* | X | + | + | + |
| *daf-19* | II | - | + | + |
| *attf-5* | X | - | + | + |
| *somi-1* | V | - | + | +/- |
| *flh-2* | III | - | - | - |
| *gfi-3* | X | - | - | - |
| *nhr-48* | X | - | - | - |
| Chromatin |  |  |  |  |
| *jmjd-3.1* | X | - | - | - |
| *utx-1* | X | - | - | - |
| Phospho. |  |  |  |  |
| *pek-1* | X | - | + | + |
| *kgb-1* | IV | + | - | - |
| Others |  |  |  |  |
| *avr-14* | I | + | + | + |
| *crb-1* | X | + | + | + |
| *aldo-1* | III | - | + | + |
| *fhod-1* | I | - | - | + |
| *puf-9* | X | - | + | - |
| *miz-1* | IV | - | - | - |
| *ncam-1* | X | - | - | - |
| TGF beta |  |  |  |  |
| *daf-5* | II | + | + | + |
| *kin-29* | X | + | + | + |
| *obr-3* | X | + | +/- | + |
| *sma-3* | III | - | + | + |
| *sma-9* | X | - | + | + |
| *sma-10* | IV | - | + | + |

Table S5: RNAi screening for delay of the onset of sterility in a *set-2*(*lf*) background. Number of progeny was estimated in F4, F8 and F12 generations. ‘-‘ indicates that number of progeny is equivalent to the one observed with the L4440 control food. ‘+’ indicates that number of progeny is greater to what is observed with the L4440 control food.
